# Characterization of a uranium-tolerant green microalga of the genus *Coelastrella* with high potential for the remediation of metal-polluted waters

**DOI:** 10.1101/2023.06.29.546994

**Authors:** Camille Beaulier, Marie Dannay, Fabienne Devime, Célia Baggio, Nabila El Sakkout, Camille Raillon, Olivier Courson, Jacques Bourguignon, Claude Alban, Stéphane Ravanel

## Abstract

Uranium (U) pollution of terrestrial and aquatic ecosystems poses a significant threat to the environment and human health because this radionuclide is chemotoxic. Characterization of organisms that tolerate and accumulate U is critical to decipher the mechanisms evolved to cope with the radionuclide and to propose new effective strategies for bioremediation of U-contaminated environments. Here, we isolated a unicellular green microalga of the genus *Coelastrella* from U-contaminated wastewater. We showed that *Coelastrella* sp. PCV is much more tolerant to U than *Chlamydomonas reinhardtii* and *Chlorella vulgaris*. *Coelastrella* is able to accumulate U very rapidly, then gradually release it into the medium, behaving as an excluder to limit the toxic effects of U. The ability of *Coelastrella* to accumulate U is remarkably high, with up to 600 mg U sorbed per g dry biomass. *Coelastrella* is able to grow and maintain high photosynthesis in natural metal-contaminated waters from a wetland near a reclaimed U mine. Over a single one-week growth cycle, *Coelastrella* is able to capture 25-55% of U from contaminated waters and demonstrates lipid droplet accumulation. *Coelastrella* sp. PCV is a very promising microalga for the remediation of polluted waters with valorization of algal biomass that accumulates lipids.

**Graphical abstract:** 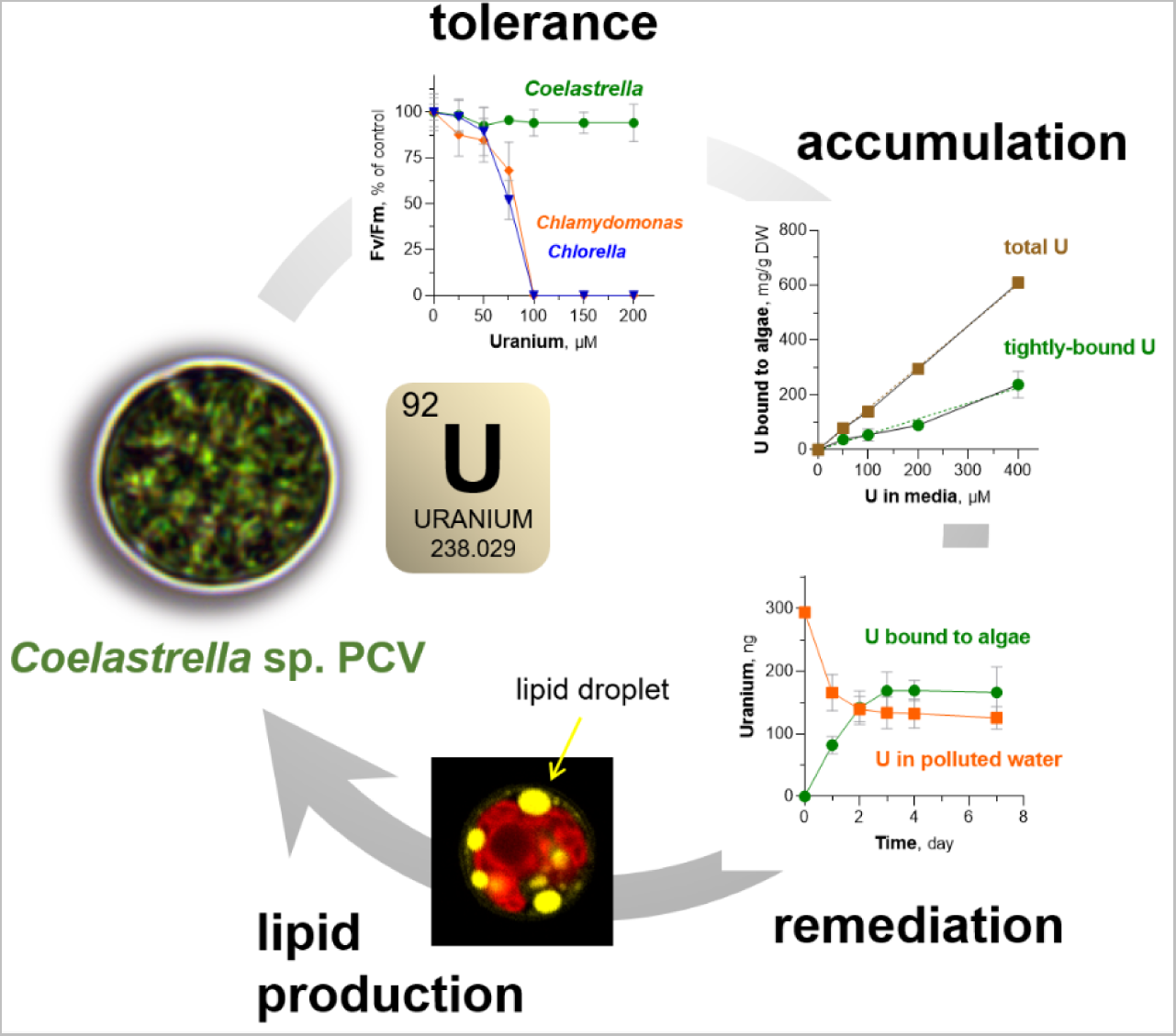

## 1. Introduction

Over the last several decades, the anthropogenic pollution of aquatic ecosystems has emerged as a major issue with global implications (Hader 2020). Freshwater ecosystems cover less than 1% of the Earth’s surface but are important reservoirs of biodiversity. Consequently, their pollution not only compromises the supply of drinking water but also poses a threat to many organisms. One of the main source of ecosystem pollution is trace metal elements, also known as heavy metals. These elements are of particular concern because they are not degradable, persistent, and hazardous to all living organisms. Conventional physical and chemical technologies for metal remediation are costly, poorly effective in treating large metal-contaminated water bodies, and can have adverse effects on aquatic ecosystems. Phytoremediation is an emerging, ecofriendly, sustainable and cost-effective green technology using metal-accumulating land or aquatic plants, macro- or micro-algae, to remove toxic metals from polluted soils and waters (Plohn 2021a; Bhat 2022). Microalgae are also promising organisms to combine wastewater reclamation and biomass valorization, as they accumulate high-value compounds such as neutral lipids and carotenoids in response to stress (Nayana 2022). In this context, the characterization of organisms with high metal tolerance and accumulation capacity is particularly relevant and timely.

Uranium (U) is a ubiquitous element in the Earth’s crust, with average concentrations of 3 ppm in soils and aquatic sediments, 3 ppb in sea water, and 0.01-30 ppb in freshwater, with a strong influence of the geochemical environment (Markich 2002). This radionuclide is primarily redistributed in the environment by anthropogenic activities related to U mining and milling industries, civil and military nuclear activities, and agricultural practices, mainly through phosphate fertilizers that are significantly contaminated with U (Anke 2009; Vandenhove 2002). Natural U is composed of >99% ^238^U, an isotope that is poorly radioactive but chemically toxic to all living organisms (Gao 2019). In recent years, our knowledge of U toxicity and the molecular mechanisms used by land plants to respond to U stress has increased significantly (Chen 2021). Very recent studies shed light on the uptake of U through calcium channels in root cells (Sarthou 2022) or by endocytosis in cultured plant cells (John 2022), and identified uranium-binding proteins in *Arabidopsis thaliana* (Vallet 2023). The fate of U is less well understood in microalgae than in land plants.

Uranium toxicity has been studied in various microalgae grown in synthetic media, natural waters, or synthetic media mimicking natural waters (Franklin 2000; Hogan 2005; Gunter 2008). Uranium uptake and toxicity in microalgae depends on its speciation, as in other organisms (Markich 2002; Gao 2019; Chen 2021). Thus, U toxicity is dependent on pH (Franklin 2000; Lavoie 2014), water hardness (Charles 2002; Fortin 2007), and its bioavailability is significantly decreased in the presence of dissolved organic carbon (Trenfield 2011; Trenfield 2012; Hogan 2015). The consequences of U intoxication on microalgal cells have been mainly reported in *Chlamydomonas* and *Chlorella* species. They include a significant impairment of cell division (Franklin 2000; Charles 2002; Pradines 2005; Hogan 2005; Lavoie 2014; Trenfield 2011) and an inhibition of photosynthesis (Pradines 2005; Herlory 2013).

Most microalgae have the ability to bind high amounts of metals, and thus the potential to remove pollutants from wastewater (Plohn 2021a). The rapid, reversible, and metabolism-independent binding of metals onto living or dead algal cells is referred to as adsorption or biosorption. The incorporation of metals inside cells through active or passive transport mechanisms is referred to absorption or uptake. Several studies have described the biosorption of U onto microalgae. This process is dependent on U speciation, pH and cell viability (dead or living cells), but is independent on temperature, light or metabolic activity (Horikoshi 1979; Zhang 1997; Gunter 2008; Vogel 2010). Cell wall composition and architecture are critical determinants of metal biosorption capacity in microalgae (Spain 2021). The combination of spectroscopic and imaging methods has indicated that carboxyl, amino and organic/inorganic phosphate groups are involved in U complexation to algal cell walls (Gunter 2008; Vogel 2010; Zhang 2022). The maximum biosorption capacity for U has been investigated in various microalgae by using very diverse experimental conditions (e.g. cell density, U concentration, incubation medium). The capacity measured for living cells ranged from 4 to 75 mg of U per g of dry biomass (Horikoshi 1979; Zhang 1997; Vogel 2010; Baselga-Cervera 2018; Zhang 2022). Only a few reports described the accumulation of U inside microalgal cells. However, these results might be interpreted with caution because rigorous discrimination between adsorption and absorption is missing or cell intactness is not established (Fortin 2004; Fortin 2007; Lavoie 2014; Cheng 2023). In an extremotolerant *Chlamydomonas* sp., biosorption onto the cell wall accounted for >90% U accumulated by algal cells, with the rest irreversibly sequestrated (Baselga-Cervera 2018). Electron microscopy dispersive X-ray spectroscopy was in agreement with the presence of U deposits in both the cell wall and cytoplasm (Garcia-Balboa 2013).

Most of the information on U accumulation and toxicity in microalgae was obtained from strains that had not been previously exposed to the radionuclide. However, examination of phytoplankton diversity in former U mining and milling sites in Salamanca, Spain, indicated that a few microalgal species were able to rapidly adapt and colonize extremely U-polluted water bodies (Garcia-Balboa 2013; Baselga-Cervera 2020). The extremotolerant *Chlamydomonas* sp. isolated from this extreme environment is a very promising strain for sustainable management of U tailings water and for deciphering the genetic basis of adaptation to U stress (Baselga-Cervera 2018).

In this study, we report the isolation and characterization of a new green microalga of the genus *Coelastrella* from laboratory liquid waste contaminated with U. We showed that this species, named *Coelastrella* sp. PCV, is much more tolerant to U than the reference green microalgae *Chlamydomonas reinhardtii* and *Chlorella vulgaris*. In addition, it is able to capture remarkably high amounts of this radionuclide from artificially contaminated media, accounting for up to 60% of its dry biomass. *Coelastrella* sp. PCV was also challenged with natural waters from a reclaimed U mine, deprived of essential nutrients and contaminated with toxic elements. *Coelastrella* sp. PCV was able to grow in natural waters and to capture 25 to 55% of the U initially present. The production of lipid droplets by stressed algal cells indicated that *Coelastrella* sp. PCV is a good candidate for both the remediation of polluted waters and valorization of the algal biomass accumulating lipids.

## 2. Materials and methods

### 2.1. Algal strains and culture conditions

*Coelastrella* sp. PCV was isolated from wastes of culture media used to study U toxicity in plants (Berthet, 2018). Green algae blooms were observed in liquid wastes containing approx. 5 µM uranyl nitrate, after a few weeks of storage in daylight at room temperature. To get rid of bacterial and fungal contamination, samples were plated onto Tris-acetate-phosphate (TAP) agar solidified medium (Harris 2009) supplemented with a cocktail of antibiotics (ampicillin 500 µg.mL^−1^, cefotaxime 100 µg.mL^−1^) and a broad-spectrum fungicide (carbendazim 40 µg.mL^−1^) (Kan 2010) Two rounds of antibiotic selection were sufficient to obtain axenic cultures of *Coelastrella* sp. PCV. The strains *Chlorella vulgaris* CCAP 211/11B and *Chlamydomonas reinhardtii* CC-125 have been used in this study.

Algal strains were maintained by weekly sub-culturing in TAP medium and growth at 21°C in continuous light at 40 µmol photons s^−1^.m^−2^. For testing U toxicity, the amount of phosphate was decreased to limit the formation of precipitates (Markich 2002). The resulting TAP LoP (low phosphate; pH 7.0) medium contained 50 µM phosphate instead of 1 mM in TAP. TAP LoP was supplemented with uranyl nitrate and inoculated with algal cells from an exponentially growing culture in TAP. The growth of *Coelastrella* sp. PCV was also analyzed in photoautotrophic conditions using the Sueoka’s high salt medium (HSM) medium (Harris 2009) and natural waters from the Rophin wetland (see below). Growth rates were assessed by cell counting using a Luna automated cell counter (Logos biosystems) for both *Coelastrella* sp. PCV and *Chlorella vulgaris*, or a Countess automated cell counter (ThermoFisher) for *Chlamydomonas reinhardtii*.

All experiments based on microalgal cultures in the presence of U or in natural waters from Rophin were repeated several times independently. An independent experiment is defined as a repetition performed at a different period with n=1-3 replicates corresponding to independent cultures (flasks).

### 2.2. Water sampling at the Rophin mining site

Water samples were collected close to the former U mining site of Rophin (Puy-de-Dome, France) (Martin 2020). The Rophin site is a pilot area for a French multidisciplinary scientific network studying territories concerned by the presence of natural and anthropogenically-enhanced radioactivity (ZATU, Zone Atelier Territoires Uranifères; https://zatu.org/). Water samples were collected in June 2019 at three points of the Rophin area. Sample E1 was from circulating water collected at the southern boundary of the mine (geographic coordinates 46.00846,3.55247; measured gamma dose rate 240 nSv/h). Water samples H1 and H2 were collected in the wetland located 200 meters downstream of the Rophin site (South-West direction). Sample H2 was collected from a stream (46.0069,3.54995; 600-800 nSv/h) while sample H1 was pore water from water-saturated soil (46.00696,3.55017; 1000-1300 nSv/h). Water samples were filtrated using fritted glass within 8 hours after sampling, then stored at 4°C. They were filtrated using 0.2 µm Nylon filters before use for chemical analysis or algal cultivation assays.

### 2.3. Phylogenetic studies

Single colonies of *Coelastrella* sp. PCV picked from TAP-agar were used for PCR amplification of the 18S rDNA gene. Colonies were dispersed in sterile water, sonicated for 1 min, and the suspension was used as a DNA template in PCR reactions. Primers 5’-ACCTGGTTGATCCTGCAG-3’ and 5’-TGATCCTCCYGCAGGTTCAC-3’ were used to amplify the 18S rDNA region (Khaw 2020). PCR products were cloned in the pSC101 vector and sequenced. The partial 18S rRNA gene sequence from *Coelastrella* sp. PCV was deposited to the GenBank database with accession number OQ202154. This sequence was compared to the Chlorophyceae sequence dataset using BLASTn from NCBI (https://blast.ncbi.nlm.nih.gov/). Thirty-one 18S rDNA sequences from the top ranking list were retrieved and analyzed using the Mega X software (Kumar 2018). Sequences were aligned using Muscle (default settings). The phylogenetic analysis was done by using the Maximum Likelihood method and Kimura 2-parameter mode. A discrete Gamma distribution was used to model evolutionary rate differences among sites (5 categories (+G, parameter = 0.0500)). The rate variation model allowed for some sites to be evolutionarily invariable ([+I], 48.22% sites). The bootstrap consensus tree was inferred from 1000 replicates.

### 2.4. Photosynthetic parameters

Algal cells (200 µL) were placed in a black-bottom 96-well plate and dark-adapted for 30 min at room temperature (21°C) before measurements. Chlorophyll fluorescence was measured using a Maxi-Imaging PAM fluorometer (Heinz Walz). The maximum quantum efficiency of photosystem II photochemistry (F_v_/F_m_) was calculated as F_v_/F_m_=(F_m_-F_o_)/F_m_, where F_0_ is the minimal fluorescence in the dark-adapted state and F_m_ the maximum fluorescence in the dark-adapted state (Maxwell 2000). The electron transport rate (ETR) was calculated as ETR = Y(II) x 0.5 x 0.84 x PAR, where Y(II) is the effective quantum yield (F_m_’-F)/F_m_’ and PAR is the incident photosynthetically active radiation (pulsing from 0 to 700 µmol photons.s^−1^.m^−2^).

### 2.5. Inductively coupled plasma mass spectrometry analyses

*Coelastrella* cells challenged with U in TAP LoP medium or in natural waters were sampled every day (2 mL of culture) and centrifuged for 4 min at 9,400 *g*. Supernatants were collected and acidified with 0.65 % (w/v) nitric acid before ICP-MS analysis. Cell pellets were washed three times with 500 µL sodium carbonate (Na_2_CO_3_) 10 mM, then dried overnight at 80°C. Dried samples were resuspended in 100 µL of 65% (w/v) HNO_3_ and mineralized for 2 hours at 90°C. Samples were diluted to the appropriate concentration and analyzed using an iCAP RQ quadrupole mass instrument (Thermo Fisher Scientific GmbH, Germany). ^238^U concentration (in ppb) was determined using a standard curve and corrected using the internal standard ^172^Yb (standard acquisition mode).

For elemental analysis of waters from the Rophin area, samples were filtrated through 0.2 µm nylon filters and acidified with 0.65 % (w/v) HNO_3_. Elements were analyzed using the standard mode (^7^Li, ^11^B, ^23^Na, ^24^Mg, ^25^Mg, ^27^Al, ^31^P, ^39^K, ^95^Mo, ^98^Mo, ^107^Ag, ^109^Ag, ^111^Cd, ^206^Pb, ^208^Pb, ^238^U) and the collision mode with helium as a cell gas (^31^P, ^39^K, ^44^Ca, ^52^Cr, ^53^Cr, ^55^Mn, ^56^Fe, ^57^Fe, ^59^Co, ^63^Cu, ^65^Cu, ^64^Zn, ^66^Zn, ^69^Ga). Quantification was as described above using the internal standards ^103^Rh and ^172^Yb.

### 2.6. Light microscopy and Nile Red staining of lipid droplets

The morphological analysis of *Coelastrella* cells was done using a Zeiss Axioplan 2 microscope. Images were acquired in bright field with magnification x63/x100 and post-processed using Fiji/ImageJ. The accumulation of lipid droplets was monitored using the Nile Red fluorescent staining procedure (Jaussaud 2020). *Coelastrella* cells grown in HSM and waters from the Rophin area were incubated for 20 min in darkness with Nile Red (Sigma Aldrich) at a concentration of 0.5 µg.mL^−1^. Cells were imaged using an inverted confocal microscope (LSM880 Zeiss) equipped with a 63x/1.4NA oil objective. Laser beams at 561 nm and 633 nm were used to excite Nile Red and chlorophyll, respectively. The software control was done using Zen Black and image post-processing was done using Fiji/ImageJ.

### 2.7. Determination of total dissolved organic carbon

Water samples filtrated using 0.2 µm Nylon filters were used for carbon analysis. Organic carbon concentration was determined by catalytic combustion (680°C) using a TOC-VCPH analyzer (Shimadzu). Total dissolved organic carbon (TOC) was determined by two methods. First, TOC was calculated from inorganic carbon (IC) and total carbon (TC) measurements using the formula TOC=TC-IC. Second, non-purgeable organic carbon (NPOC) measurements were carried out to verify TOC values obtained by the TC-IC determination.

## 3. Results

### 3.1. Isolation, genetic identification, growth and morphological characteristics of *Coelastrella* sp. PCV

The development of a green algal bloom was observed in laboratory liquid wastes from culture media used for the study of U stress in plants (Berthet 2018). In order to identify the green alga(e) able to grow in this medium contaminated by uranyl nitrate (UO_2_(NO_3_)_2_); about 5 µM, 1.2 ppm), we performed two rounds of selection on antibiotics/fungicide and obtained an axenic algal culture (**Figure S1**). Colony PCR with 18S rDNA primers was used for the genetic identification of selected microalgae. A unique partial 18S rDNA sequence was obtained from all tested colonies. It showed >99% sequence identity with 18S rDNA sequences of microalgae from the Scenedesmaceae family (within the Chlorophyceae class and Sphaeropleales order), including members of the genera *Coelastrella*, *Scenedesmus*, *Tetradesmus* or *Asterarcys*. A phylogenic tree inferred from 31 related 18S rDNA sequences showed that the isolated microalga was clustered with *Coelastrella* species (**Figure 1A**). The genus *Coelastrella* was first reported by Chodat (Chodat 1922) and 24 species isolated from aero-terrestrial or freshwater habitats are currently described in the AlgaeBase (algaebase.org) (Guiry 2021). The strain selected from liquid wastes probably originates from airborne contamination of the laboratory environment. Since the isolate does not correspond to any described species, it was named *Coelastrella* sp. PCV (with reference to the laboratory acronym).

**Figure 1:**
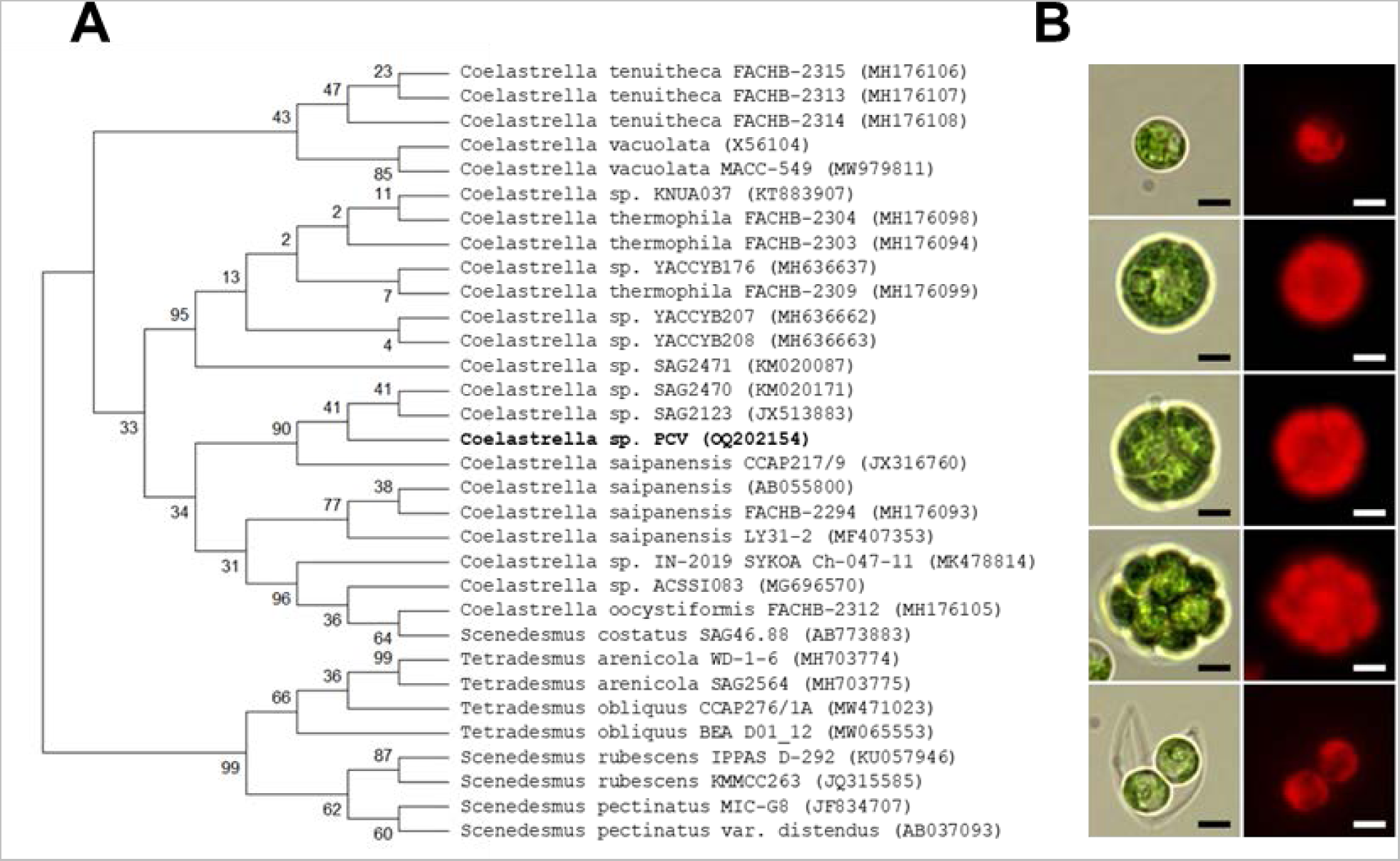
Phylogenetic and morphological analysis of *Coelastrella* sp. PCV. A-Phylogenetic analysis inferred from 18S rDNA sequences. The 18S rDNA sequence from *Coelastrella* sp. PCV was obtained by PCR and used for a NCBI BLASTn search against the Chlorophyceae sequence dataset. Thirty-one 18S rDNA sequences from the top ranking list were retrieved and aligned using Muscle (MEGA X software, default settings). The phylogenetic analysis was done by using the Maximum Likelihood method and Kimura 2-parameter mode. The consensus bootstrap tree is shown and branch support values (in % for 1000 replicates) are indicated. The GenBank accession numbers are shown in brackets. B-Morphological analysis of *Coelastrella* sp. PCV at different stages of growth. Bright-field and chlorophyll fluorescence pictures were imaged using a Zeiss Axioplan 2 microscope and post-processed using ImageJ/Fiji. Scale bar = 5 µm.

*Coelastrella* sp. PCV was maintained under mixotrophic growth conditions in Tris-acetate-phosphate (TAP) medium in continuous light, either in liquid cultures with weekly subcultures or on TAP-agar plates. The microalga was also able to grow under photoautotrophic conditions in media such as the Sueoka’s high salt medium (HSM) or Bold’s basal medium (BBM), in continuous light or light/dark cycles. *Coelastrella* sp. PCV is a fast-growing microalga in mixotrophic or autotrophic conditions and batch cultures reach a maximal cell density of 5-6 million cells.mL^−1^ at the stationary phase. The maximal cell density is quite low and can be explained by the large size of algal cells (Agusti 1989). Indeed, the morphological analysis of *Coelastrella* sp. PCV by light microscopy showed green unicellular and spherical cells with an average diameter of 9 ± 3 µm (**Figure 1B**). Cell size distribution in liquid cultures is quite variable, with young cells of 7 ± 1 µm in diameter and mature cells up to 16 ± 2 µm in diameter. Vegetative reproduction occurs by autosporulation, with sporangia containing 2 to 16 daughter cells (**Figure 1B**). Autospores are released after the rupture of the parental cell wall. Sexual reproduction was not observed in standard growth conditions.

### 3.2. Tolerance of *Coelastrella* sp. PCV to uranium in synthetic media

The isolation of *Coelastrella* sp. PCV from liquid wastes contaminated by U prompted us to analyze its tolerance to this radionuclide that is chemically toxic to photosynthetic organisms (Chen 2021). The speciation of U in natural or artificial environments is very complex and determines its toxicity to living organisms (Gao 2019). In particular, the presence of phosphate in plant or algal culture media is known to limit U bioavailability due to U-phosphate complex formation and precipitation (Markich 2002; Fortin 2004; Pradines 2005). We found that phosphate limitation in TAP medium did not impaired significantly the growth of *Coelastrella* sp. PCV (**Figure 2**). In contrast, phosphate limitation in phototrophic media was accompanied with a drastic reduction in cell division, which was not compatible for measuring U toxicity. Thus, the tolerance of *Coelastrella* sp. PCV to U stress was analyzed in a TAP medium containing 50 µM of phosphate (TAP LoP).

**Figure 2:**
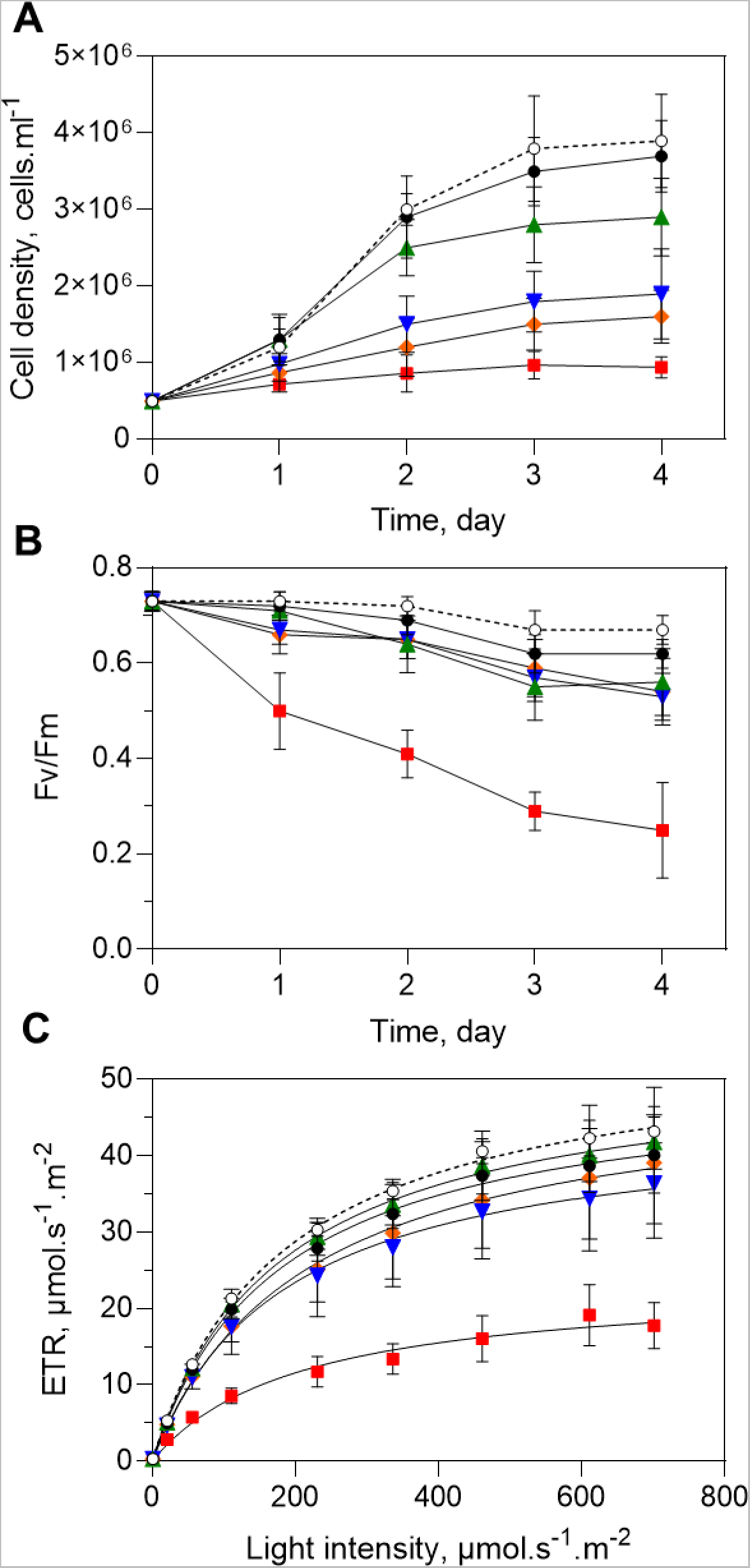
Effect of uranium on the growth and photosynthesis of *Coelastrella* sp. PCV. A-Growth curves were established for cells growing in TAP (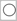) or TAP LoP (50 µM of phosphate) supplemented with uranyl nitrate (0 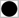, 50 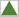, 100 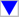, 200 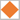, and 400 µM 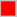). Cells were grown at 21°C in continuous white light (40 µE) and counted daily. B,C-Photosynthetic efficiency of *Coelastrella* cells challenged with U. Fv/Fm in dark-adapted cells (B) and electron transfer rates (ETR) under increasing light irradiance were determined after 2 days of U stress. Data from n=5 to 8 independent experiments (with n=2-3 replicates) analyzed over a 3-year period have been gathered to produce growth and Fv/Fm kinetics (mean ± SD). Light response curves are representative of n=2 independent experiments with n=3 replicates (mean ± SD).

*Coelastrella* sp. PCV cells were challenged with 50 to 400 µM uranyl nitrate in TAP LoP for 4 days. Growth was measured by cell counting and photosynthetic activity was used as a proxy to assess U toxicity (Herlory 2013). As shown in **Figure 2A**, algal growth was inhibited by U in a dose-dependent manner. At the highest U concentration tested (400 µM), the growth of *Coelastrella* cells was severely impaired. Determination of the doubling time during the exponential growth phase allowed calculation of a half-maximal inhibitory concentration (IC_50_) of 80 µM **(Figure S2A)**. Chlorophyll fluorescence was used to measure the maximum quantum yield of photosystem II in dark-adapted cells (F_v_/F_m_) and the electron transport rate (ETR) at various light intensities (Maxwell 2000). We did not observed significant effect of U on photosynthetic parameters for concentrations up to 200 µM. However, a significant decrease in Fv/Fm was observed at 400 µM uranyl nitrate, starting from 1 day of treatment and reaching 65% reduction after 4 days of U exposure (**Figure 2B**). In addition, the ETR measured during the exponential growth phase was decreased by 40% at the highest light intensity (700 µmol photons.s^−1^.m^−2^) compared to the control TAP LoP condition **(Figure 2C)**. Dose-effect relationships gave IC50 values of >400 µM for ETR and >500 µM for F_v_/F_m_ (**Figure S2B-C**), indicating that the photosynthetic machinery of *Coelastrella* cells was moderately affected by U stress. In line with these results, when cells challenged for 4 days with 400 µM uranyl nitrate were reintroduced into fresh TAP medium, growth and photosynthetic rates were comparable to unstressed cells, demonstrating that *Coelastrella* sp. PCV was able to survive severe U stress. Together, these experiments showed that *Coelastrella* sp. PCV is tolerant to high U concentrations. Noteworthy, tolerance experiments have been conducted over a 3-year period since the isolation of the microalga. Growth curves and photosynthetic properties were similar throughout the period, indicating that the ability of *Coelastrella* sp. PCV to tolerate U is genetically fixed.

The physiological consequences of U stress in green microalgae have been reported in the models *Chlamydomonas* and *Chlorella* (Franklin 2000; Charles 2002; Pradines 2005; Hogan 2005; Trenfield 2011; Herlory 2013; Lavoie 2014). Comparing U tolerance of *Coelastrella* sp. PCV (this work) with data from the literature is not relevant as U speciation, bioavailability and therefore toxicity differ significantly due to the variety of stress conditions. Therefore, we included *Chlamydomonas reinhardtii* and *Chlorella vulgaris* in our study for comparison. As previously, cell growth and photosynthetic parameters were measured to assess tolerance to U stress in TAP LoP containing uranyl nitrate. The inhibitory effect of U in *C. reinhardtii* and *C. vulgaris* was observed from 50 µM uranyl nitrate (**Figure 3**), and at 100 µM growth was fully abolished and Fv/Fm values were null, indicating that the cells were dead. Under these conditions, the growth of *Coelastrella* sp. PCV was only reduced by 50% and F_v_/F_m_ was not affected. These data show that *Coelastrella* sp. PCV is much more tolerant to U than freshwater model microalgae.

**Figure 3:**
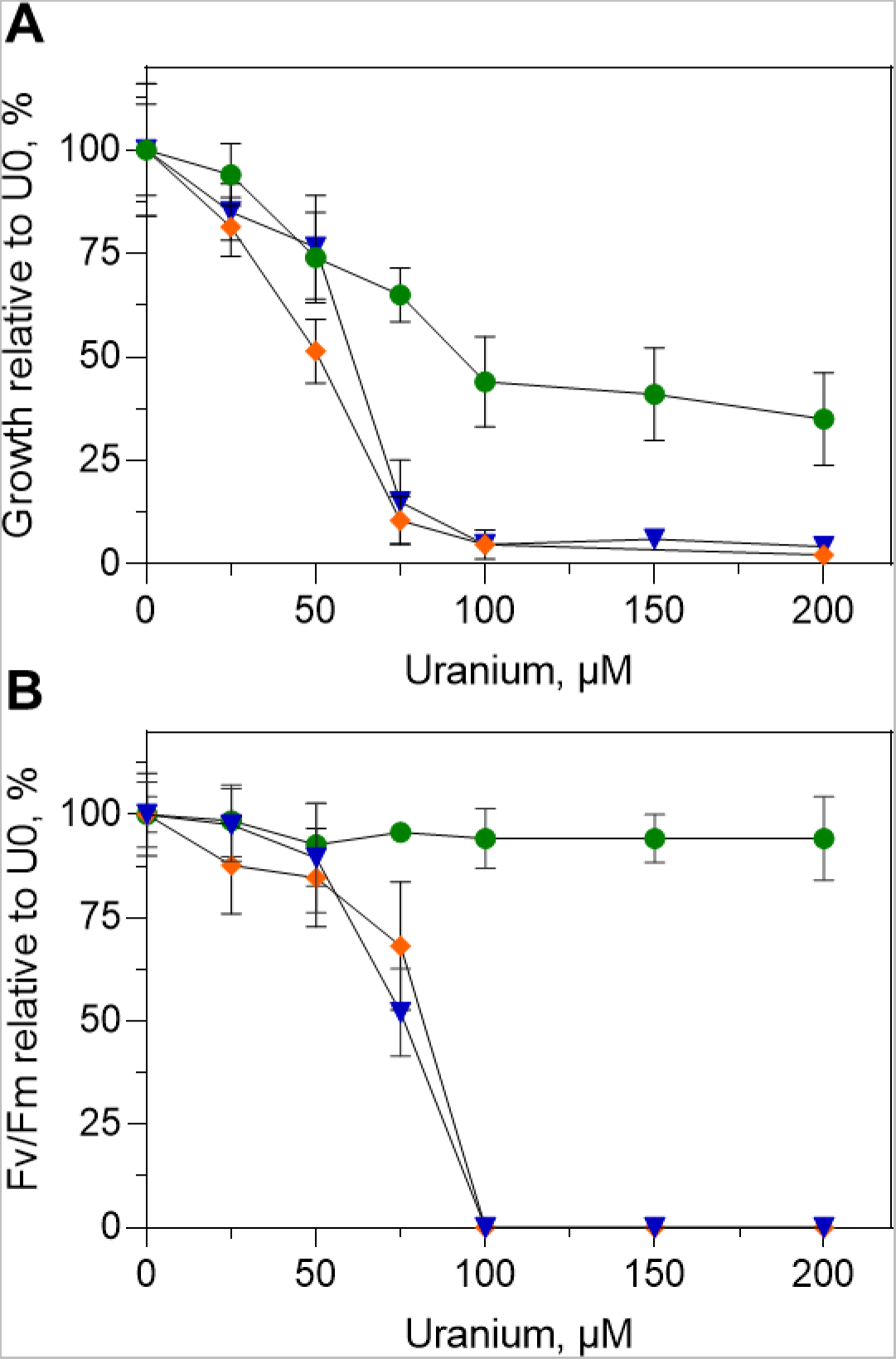
Comparison of uranium tolerance in *Coelastrella* sp. PCV, *Chlamydomonas reinhardtii* and *Chlorella vulgaris*. A-Growth curves were established for *Coelastrella* sp. PCV (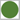), *Chlamydomonas reinhardtii* (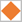) and *Chlorella vulgaris* (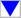) grown in TAP LoP containing 0-200 µM of uranyl nitrate. Cells were grown at 21°C in continuous white light (40 µE) and cell density after 2 days of U stress was used to calculate relative growth with reference to TAP LoP (U0). B-Fv/Fm was determined in dark-adapted cells after 2 days of U stress. Data from n=3 to 5 independent experiments (and n=2-3 replicates) are displayed (mean ± SD).

Next, we addressed the fate of U in algal cultures over a 4-day incubation period. The amount of U remaining in the medium and tightly bound to algal cells was determined by ICPMS. To this end, algal cells were extensively washed with sodium carbonate to remove U that is loosely adsorbed to the cell wall (Revel 2022; Sarthou 2022). For uranyl nitrate concentrations ranging from 50 to 200 µM, the kinetics of U accumulation in the medium and cells clearly showed two main phases **(Figure 4A-C)**. During the first 24 hours, U was markedly depleted from the medium and reached its highest accumulation level in the cells. Eighty to 95% of U initially present in the medium was captured by algal cells, of which a large amount was tightly bound and could not be released with carbonate. From 48 hours, a second phase corresponding to a gradual release of U from the cells to the medium was observed **(Figure 4A-C)**. As a result of this release process, the amount of U in the medium returned to 50 to 80% of its initial concentration after 4 days of culture. When cells were challenged with 400 µM uranyl nitrate, the first phase was similar to that observed for lower metal concentrations, with a strong decrease (90%) of U in the medium and the accumulation of tightly-bound U in the cells (**Figure 4D**). The second phase was different, however, because we did not observe U release from cells but, instead, constant levels of U in the cells and medium until the end of the stress period **(Figure 4D)**. These data suggest that U release from *Coelastrella* cells is a defense process whose efficiency is dependent on stress intensity and algal physiology (**Figure 2**).

**Figure 4:**
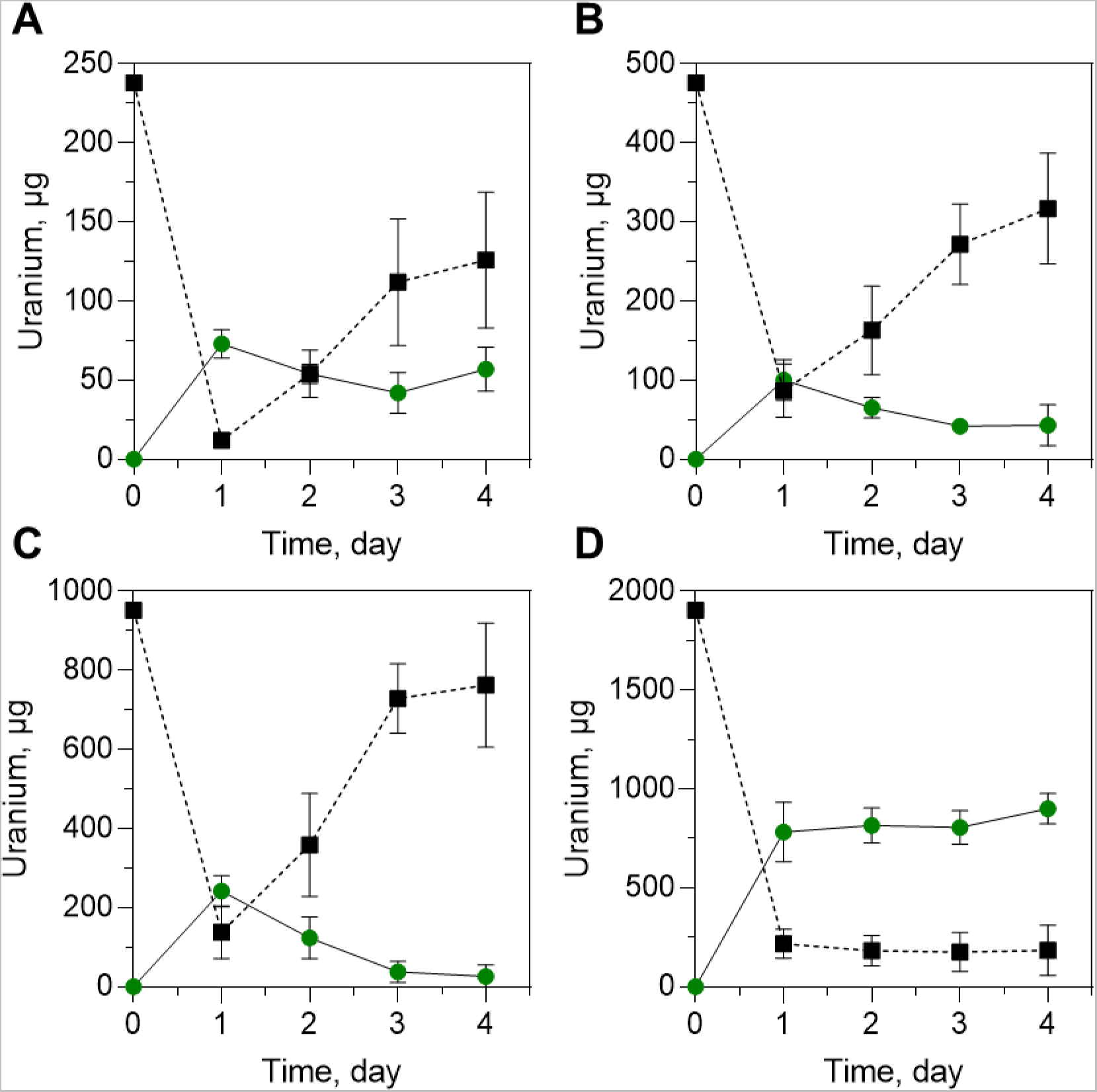
Time course of uranium distribution between the medium and algal cells during growth under stress conditions. *Coelastrella* sp. PCV was grown at 21°C in continuous white light (40 µE) in TAP LoP containing 50 (A), 100 (B), 200 (C) and 400 (D) µM of uranyl nitrate (see Figure 2). Aliquots of the medium (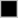) and cells (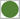) were withdrawn every day and U was determined by ICPMS. Before U quantification, cells were washed with sodium carbonate to remove loosely-bound metals. The amount of U (in µg) is indicated for a standard 20-mL culture. Data from n=5 independent experiments (and n=2-3 replicates) are shown (mean ± SD).

### 3.3. Accumulation of uranium in *Coelastrella* sp. PCV

To gain further insight into the accumulation of U by *Coelastrella* sp. PCV, we analyzed the fate of the radionuclide during the first 24-hour period. Algal cells were incubated with different concentrations of uranyl nitrate (from 50 to 400 µM) in the TAP LoP medium and aliquot fractions were withdrawn at regular time intervals to measure U in the medium, in whole cells, and in cells washed with sodium carbonate. As shown in **Figure 5A**, 80 to 98% of U initially present in the medium was depleted within 1.5 hours and the amount of U was maintained at a very low level up to 24 hours. A very rapid accumulation of U in *Coelastrella* whole cells was concomitant with these kinetics (**Figure 5B**). At 24 hours of incubation, the amount of U tightly bound to algal cells ranged from 30 to 50% of total U (**Figure 5C**). A plot of the amount of U accumulated in cells (on a dry mass basis) as function of the initial U concentration in the medium was used to estimate the maximum capacity of *Coelastrella* cells to capture U. As shown in **Figure 5D**, both the total amount of U and the tightly-bound fraction increased linearly with U concentration in the medium. These data indicated that the accumulation (adsorption and absorption) of U by *Coelastrella* cells was not saturated at 400 µM uranyl nitrate in TAP LoP medium. In these conditions, *Coelastrella* sp. PCV was able to capture up to 600 mg U per gram dry biomass, of which 240 mg.g^−1^ DW was tightly bound and/or incorporated within algal cells. Together, these data show that *Coelastrella* sp. PCV has a remarkably high capacity to accumulate U.

**Figure 5:**
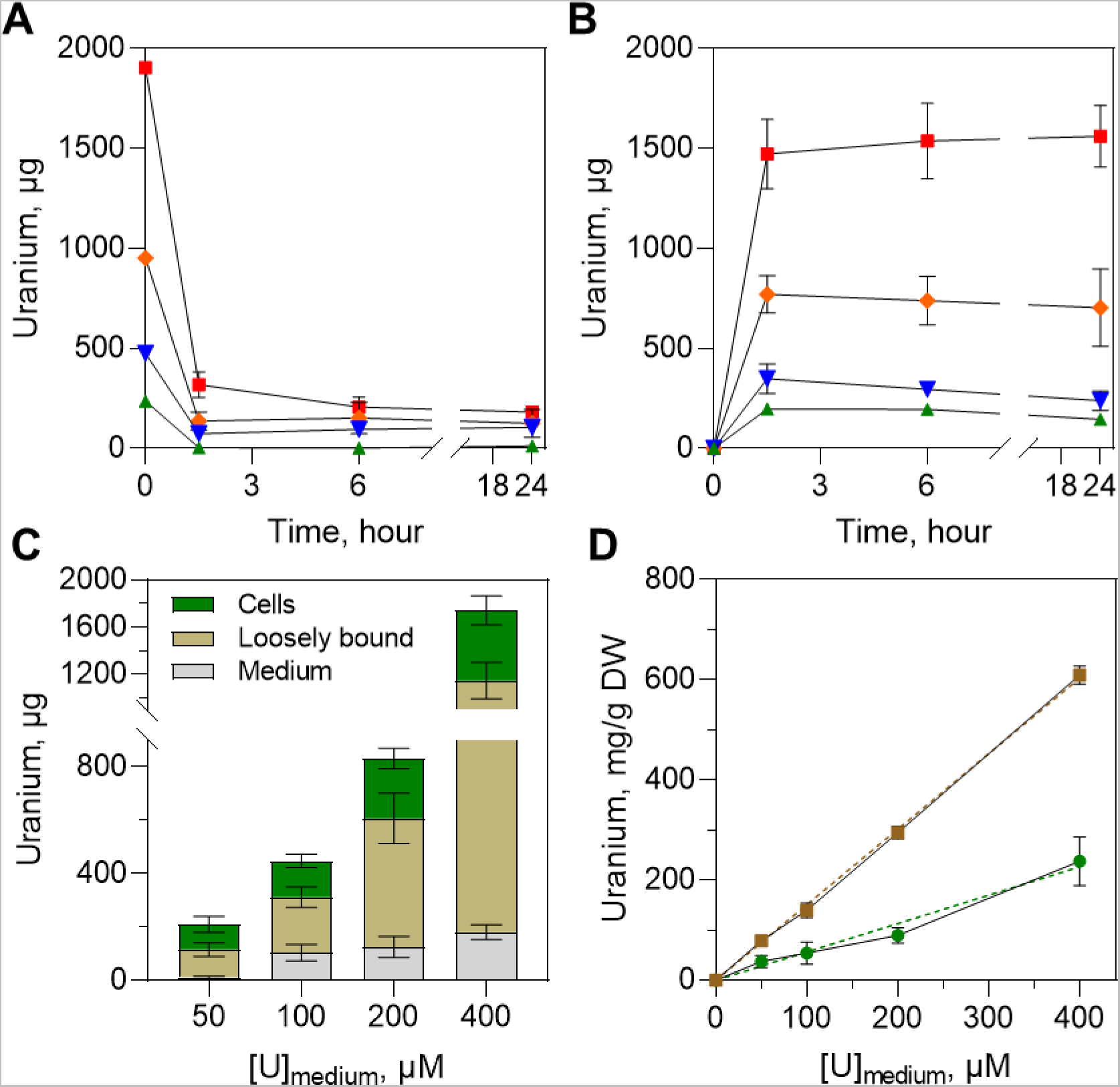
Biosorption of uranium by *Coelastrella* sp. PCV. Time course of U distribution between the medium (A) and algal cells (B) during short-term incubation periods. *Coelastrella* sp. PCV was challenged with 50 (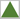), 100 (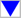), 200 (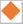) and 400 (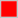) µM uranyl nitrate in TAP LoP at 21°C in continuous white light (40 µE). Uranium was determined by ICPMS in the medium and whole cells (not washed with sodium carbonate) at different time intervals. C-Quantification of loosely and tightly-bound U to *Coelastrella* cells. Cells were washed with sodium carbonate to remove loosely-bound U. Data show the distribution of U at time point 24 hours. In panels A,B,C, the amount of U (in µg) is indicated for a standard 20-mL culture. D-Titration of U biosorption capacity of *Coelastrella* cells. The amount of metal bound to whole cells (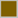) or carbonate-washed cells (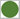) is expressed in mg U per gram dry weight. Data from n=5 independent experiments (and n=1-3 replicates) are shown (mean ± SD).

### 3.4. Remediation of natural polluted waters by *Coelastrella sp*. PCV

To analyze the ability of *Coelastrella* sp. PCV to remediate natural waters polluted by U, we collected water samples at the vicinity of the former U mining site of Rophin (Puy-de-Dome, France). Ore bodies from the Rophin site are rich in parsonsite, a lead-uranium-phosphate mineral (Pb_2_(UO_2_)(PO_4_)_2_,·2H_2_O), and were exploited between 1949 and 1957 (Martin 2020). During this period and before complete rehabilitation of the site, the wetland located a few hundred meters downstream from the mine accumulated U of anthropogenic origin. Water samples were collected from three locations of the Rophin site with measured gamma dose rates of 240, 600-800 and 1000-1300 nS.h^−1^ for samples E1, H2, and H1, respectively. The local geological background of the area was 200-250 nS.h^−1^ (Martin 2020). The elemental composition of waters from Rophin was analyzed by ICPMS (**Table 1**). Waters were found to be quite poor in elements that are essential for microalgal growth, with magnesium, potassium, phosphorus, calcium or iron in the micromolar concentration range, and manganese, copper, zinc or molybdenum in the nanomolar concentration range. Also, Rophin waters contain non-essential and potentially toxic elements, including U at a concentration of 15 ± 1, 62 ± 12 and 228 ± 40 nM in the E1, H2, and H1 waters, respectively (**Table 1**). Total dissolved organic carbon (TOC) was also determined using the differential (total carbon minus dissolved inorganic carbon) and NPOC methods. Both methods gave similar results and indicated that the H1 pore water contained 3 to 4-times more TOC than the E1 and H2 flowing waters (**Table 1**).

**Table 1:** Physico-chemical properties of natural waters from the Rophin area. Water samples from the Rophin area were collected at the E1, H1 and H2 locations and filtered using fritted glass and then 0.2 µm nylon filters before analysis. Elemental composition was determined by ICPMS. Data are mean ± SD of n=3-5 measurements. Total carbon (TC), dissolved inorganic carbon (DIC), and non-purgeable organic carbon (NPOC) were determined by the high temperature catalytic oxidation methods and non-dispersive infrared detection. Data are mean ± SD of n=3-6 measurements with n=2 technical replicates.

|  | E1 | H1 | H2 | Unit |
| --- | --- | --- | --- | --- |
| **pH** | 7.0-7.2 | 6.9-7.1 | 7.0-7.3 |  |
| **Na** | 223 ± 46 | 214 ± 27 | 214 ± 24 |  |
| **Mg** | 31 ± 0.8 | 91 ± 7.8 | 47 ± 2.9 |  |
| **Al** | 1.1 ± 0.4 | 2.2 ± 0.4 | 1.3 ± 0.4 |  |
| **P** | 0.73 ± 0.1 | 1.23 ± 0.5 | 0.80 ± 0.2 | x10<sup>-6</sup> M |
| **K** | 24 ± 1.2 | 37 ± 1.9 | 36 ± 2.1 |  |
| **Ca** | 66 ± 12 | 151 ± 41 | 97 ± 24 |  |
| **Fe** | 4.4 ± 0.8 | 53.9 ± 5.6 | 4.5 ± 1.0 |  |
| **Li** | 79 ± 22 | 106 ± 30 | 95 ± 32 |  |
| **B** | 206 ± 79 | 261 ± 102 | 510 ± 159 |  |
| **Cr** | 2.0 ± 0.3 | 11.8 ± 2.8 | 3.3 ± 0.7 |  |
| **Mn** | 70 ± 29 | 9260 ± 2909 | 69 ± 25 |  |
| **Co** | 0.8 ± 0.3 | 38.0 ± 17.8 | 1.0 ± 0.2 |  |
| **Cu** | 32 ± 8 | 327 ± 74 | 135 ± 22 |  |
| **Zn** | 89 ± 26 | 305 ± 66 | 90 ± 13 | x10<sup>-9</sup> M |
| **Ga** | 53 ± 6 | 199 ± 28 | 81 ± 8 |  |
| **Mo** | 1.5 ± 0.4 | 10.3 ± 2.8 | 1.6 ± 0.3 |  |
| **Ag** | 0.12 ± 0.06 | 0.21 ± 0.14 | 0.36 ± 0.11 |  |
| **Cd** | 0.20 ± 0.03 | 0.60 ± 0.14 | 0.40 ± 0.06 |  |
| **Pb** | 3.8 ± 0.5 | 277 ± 71 | 21 ± 8 |  |
| **U** | 15 ± 1 | 228 ± 40 | 62 ± 12 |  |
| **Total carbon (TC)** | 8.2 ± 0.5 | 24.5 ± 1.2 | 8.9 ± 0.4 |  |
| **Inorganic carbon (IC)** | 4.0 ± 0.2 | 5.8 ± 2.9 | 4.1 ± 1.1 | mg C.l<sup>-1</sup> |
| **Total organic carbon (TC – IC)** | 4.2 ± 0.6 | 18.7 ± 3.9 | 4.8 ± 1.1 |  |
| **Total organic carbon (NPOC)** | 3.8 ± 0.3 | 14.8 ± 2.8 | 4.4 ± 0.7 |  |

We analyzed the ability of *Coelastrella* sp. PCV to proliferate in Rophin waters. As shown in **Figure 6A**, *Coelastrella* sp. PCV was able to grow exponentially in all three waters to reach a maximal cell density of 2-2.5 million cells.mL^−1^. The stationary phase was reached 4 days after inoculation while growth continued in the control HSM medium, suggesting growth impairment due to nutrient limitation and/or toxicity of contaminating metals (e.g. U, lead, aluminum) (**Table 1**). Monitoring of Fv/Fm and ETR indicated that *Coelastrella* was able to maintain significant photosynthetic activity for at least 7 days of growth in Rophin waters (**Figure 6B-C**).

**Figure 6:**
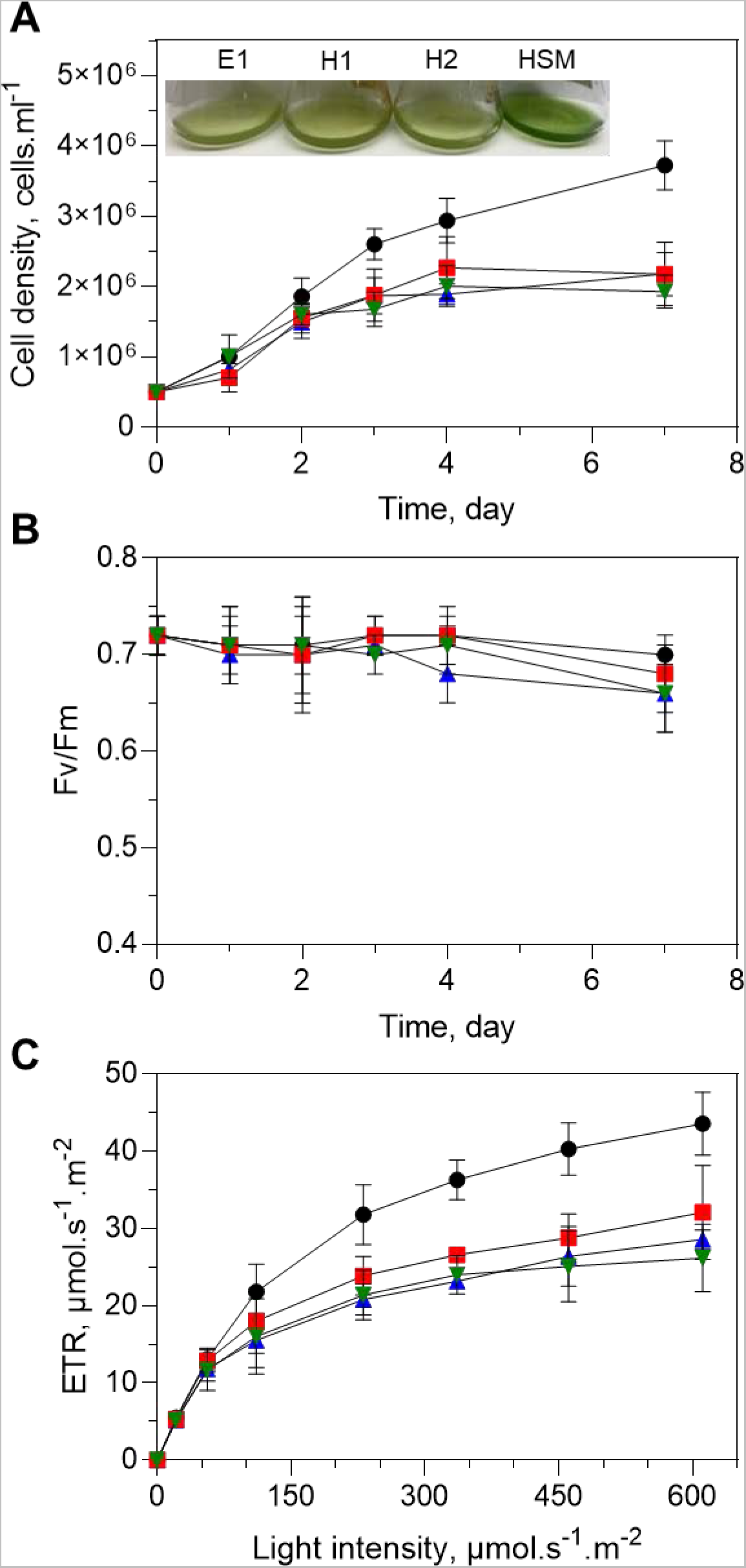
Growth and photosynthesis of *Coelastrella* sp. PCV in natural metal-polluted waters. A-Growth curves were established for cells growing in HSM (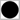) and natural waters from the Rophin area (E1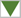, H1 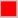, H2 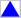). Cells were grown at 21°C in continuous white light (40 µE) and counted daily. Picture in the insert shows cultures on day 7. Data from n=5 independent experiments with n=2 replicates are shown (mean ± SD). B,C-Photosynthetic efficiency of *Coelastrella* cells grown in natural waters. Fv/Fm was determined in dark-adapted cells. Data from n=5 independent experiments with n=2 replicates are shown (mean ± SD). The ETR was determined under increasing light irradiance after 3 days of growth. Data from n=2 independent experiments with n=2 replicates are shown (mean ± SD).

The distribution of U between polluted waters and *Coelastrella* cells was analyzed during a 7-day growing period. As the goal of this experiment was to evaluate the remediation potential of *Coelastrella* sp. PCV, we first determined the total amount of U associated with the algal biomass, including loosely (carbonate washable) and tightly (unaffected by carbonate treatment) bound U. Data in **Figure 7A-B** show that growth of *Coelastrella* sp. PCV in the E1 and H2 waters resulted in the removal of 50 to 55% of U after 2 to 3 days of growth. In the H1 water, the removal of U was slower and less efficient with a 25% decrease of U concentration in 7 days (**Figure 7C**). In contrast with accumulation profiles observed in synthetic TAP LoP medium containing up to 200 µM uranyl nitrate (**Figure 4**), there was no release of U captured by algal cells during prolonged incubation periods. After 7 days of growth in Rophin waters, 40 to 50% of U was tightly bound to algal cells and the rest could be released by washing with sodium carbonate (**Figure 7D**). In addition to U, Rophin waters are significantly contaminated by lead (**Table 1**), mostly originating from the decay chain of U as demonstrated by the analysis of lead isotopy (Martin 2020). The ability of *Coelastrella* sp. PCV cells to bind lead was analyzed. After 7 day of growth, *Coelastrella* cells captured 40 to 45% of lead present in E1 and H2 waters, and 5% from the H1 water. Thus, *Coelastrella* sp. PCV was able to reclaim waters from at least two heavy metals (U and lead) resulting from the anthropogenic contamination of the Rophin wetland.

**Figure 7:**
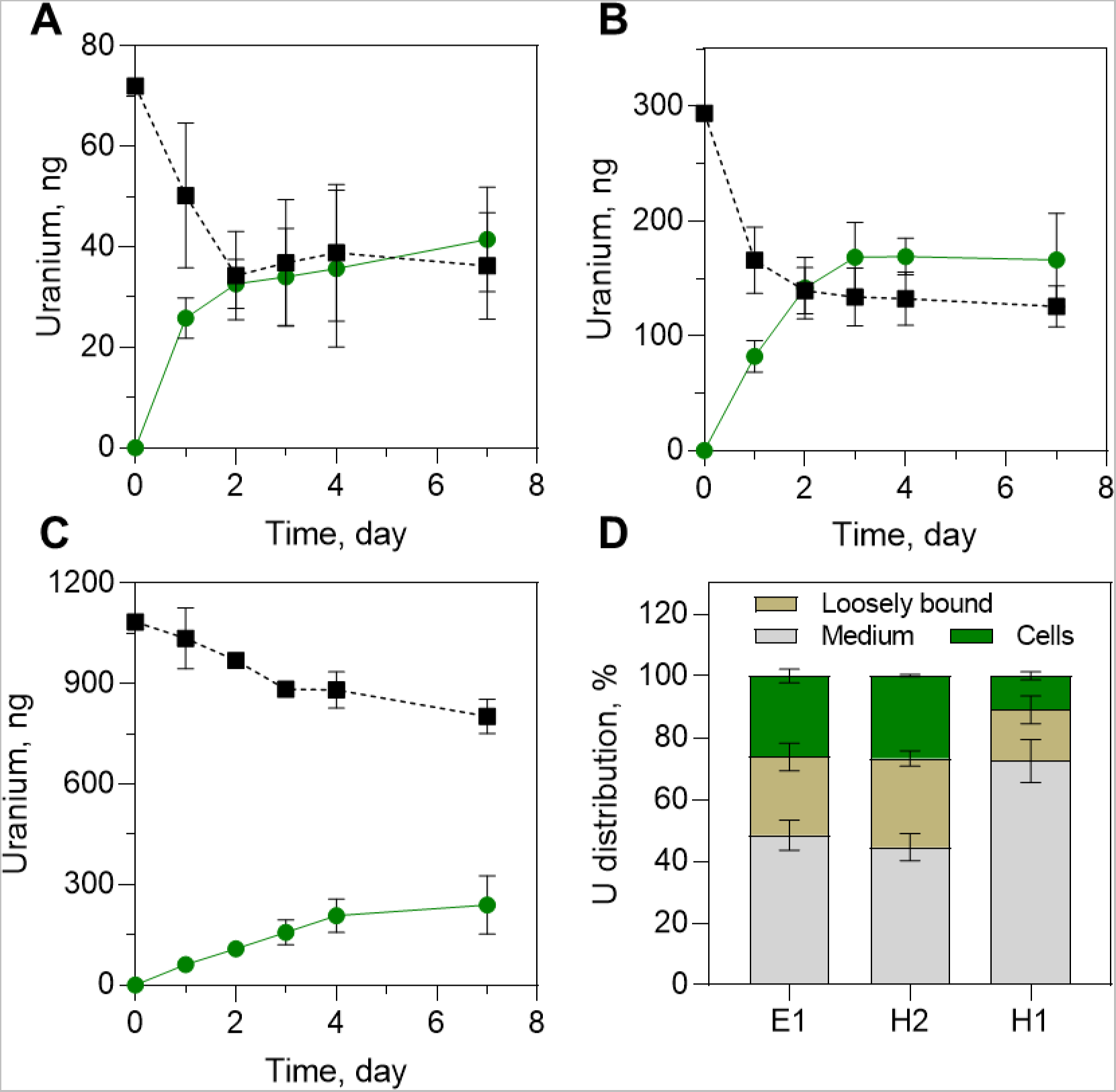
Bioremediation of uranium from natural polluted waters by *Coelastrella* sp. PCV. A,B,C-Time courses of U distribution between the medium (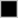) and algal cells (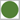) during growth of *Coelastrella* in waters from the locations E1 (A), H2 (B), and H1 (C) of the Rophin area. Growth was conducted at 21°C in continuous white light (40 µE). Uranium was determined by ICPMS in the medium and whole cells (not washed with sodium carbonate) at regular time intervals. The amount of U (in ng) is indicated for a standard 20-mL culture. D-Distribution of U remaining in waters and loosely or tightly-bound to *Coelastrella* cells. Cells were washed with sodium carbonate to remove loosely-bound U. Data show the distribution of U after 7 days of growth in contaminated waters. Data from n=3 to 6 independent experiments and n=2 replicates are shown (mean ± SD).

The growth of *Coelastrella* sp. PCV in waters from the Rophin wetland was significantly reduced as compared to a nutrient-sufficient and contaminant-free synthetic medium (**Figure 6A**). The impairment of algal cell growth due to nutrient deprivation or another stressful situation is known to trigger neutral lipid production (Sun 2018). Under adverse conditions, several *Coelastrella* species have been shown to accumulate high amount of lipids with fatty acid profiles of interest for the biofuel industry (Nayana 2022). Lipid production in *Coelastrella* sp. PCV grown in Rophin waters was investigated using the Nile red staining procedure. As shown in **Figure 8**, several large lipid droplets were observed in the cytoplasm of *Coelastrella* cells grown in either E1, H1 or H2 waters. As a comparison, cells grown in HSM displayed very small lipid droplets. These data indicate that growth of *Coelastrella* sp. PCV in nutrient-deprived and metal-polluted waters triggered a reorientation of cellular metabolism to produce and accumulate neutral lipids, representing a potential interest for industrial and technology applications.

**Figure 8:**
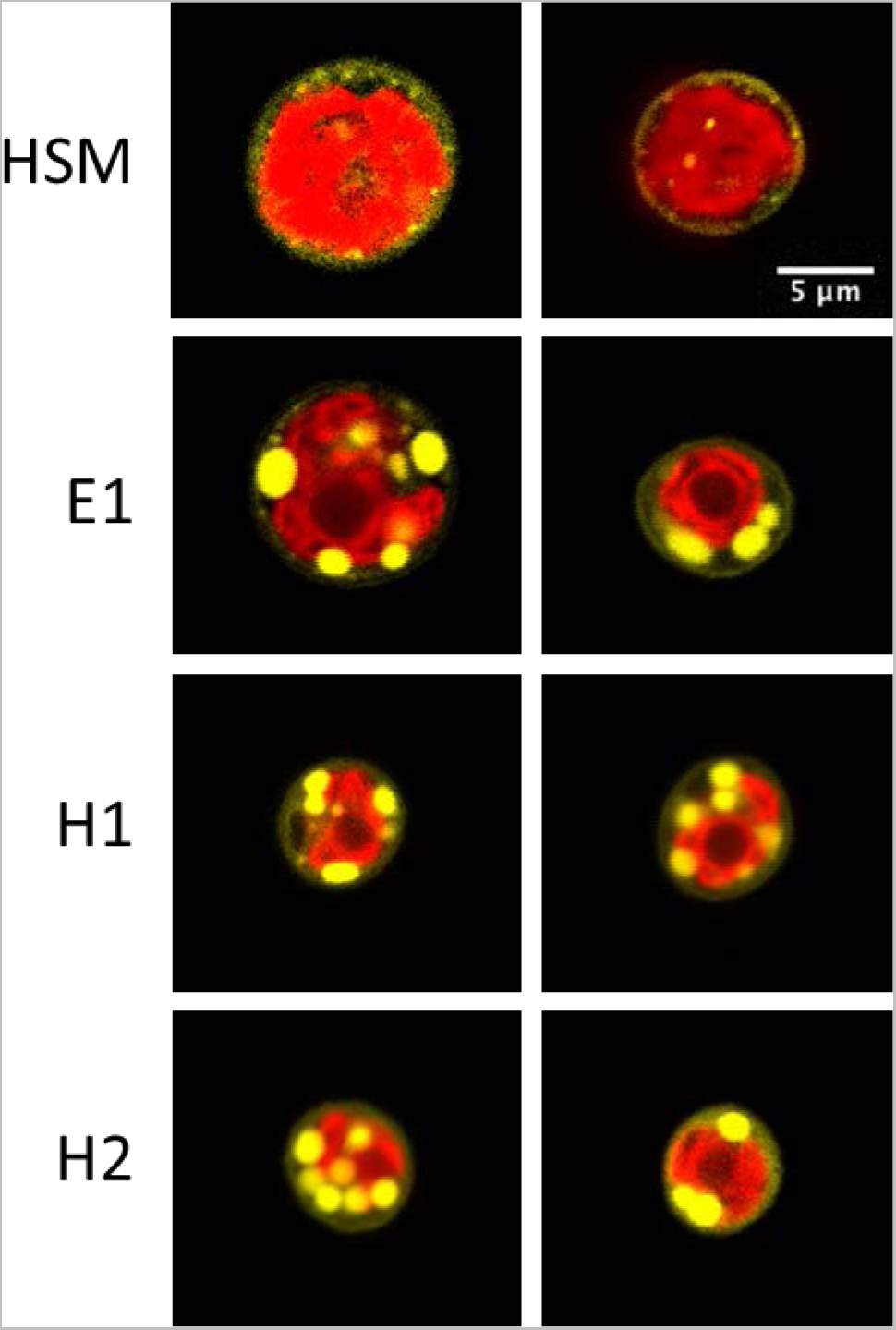
Lipid droplet accumulation in *Coelastrella* sp. PCV grown in natural polluted waters. *Coelastrella* sp. PCV was cultivated in waters from Rophin for 3 days at 21°C in continuous white light (40 µE) and then stained with Nile Red. Cells were imaged using an inverted confocal microscope (LSM880 Zeiss). Chlorophyll autofluorescence is in red; lipid droplets stained with Nile Red are in yellow. The images have been selected to highlight lipid droplets; they are not representative of cell size because the z-depth does not necessarily correspond to the median plane of cells. Additional experiments showed that the average diameter of *Coelastrella* cells grown in HSM and H1 water was 7.3 ± 2.0 and 6.0 ± 0.8 µm, respectively (mean ± SD, n=200 cells per condition).

## 4. Discussion

The green unicellular microalga isolated from U-contaminated liquid wastes belongs to the genus *Coelastrella*. Genetically, 18S rDNA sequence analysis indicated that *Coelastrella* sp. PCV is related to at least one *Coelastrella saipanensis* strain, but branch support values are not enough robust to allow accurate determination at the species level (**Figure 1A**). The morphological characteristics of *Coelastrella* sp. PCV (**Figure 1B**) did not allow to refine this classification but are in agreement with this relationship with *C. saipanensis* (Smith-Baedorf 2013; Wang 2019). The genus *Coelastrella* was first described a century ago (Chodat 1922) and is currently represented by dozen of species (Guiry 2021; Nayana 2022). These microalgae have been found worldwide in a variety of aero-terrestrial and aquatic environments. Some *Coelastrella* species are characterized by the harshness of their natural habitat, including warm water from Roman baths (Smith-Baedorf 2013), municipal wastewater in Nordic climate (Ferro 2018), or dry terrestrial surfaces exposed to high light and elevated temperature (Narayanan 2018; Kawasaki 2020).

The ability of *Coelastrella* species to tolerate metal intoxication or to live in habitats contaminated with metals has been reported. Toxicity evaluation of zinc and iron oxide nanoparticles in *Coelastrella terrestris* indicated that this species has the potential for the remediation of metallic nanoparticles (Saxena 2019; Saxena 2020). *Coelastrella* sp. (3-4) and *Coelastrella* sp. BGV displayed high absorption capacity for cadmium, copper or lead, and can remove metals from contaminated environments very efficiently (Plohn 2021b; Karcheva 2022). Also, a *Coelastrella* sp. was isolated from copper mine tailings sand (Wu 2022). Co-inoculation of this microalgal species and two native fungi onto the contaminated tailings sand resulted in a 15% decrease of copper content.

Here, we describe for the first time the physiological response and accumulation capacity of a *Coelastrella* species exposed to U. The tolerance to U has been previously investigated in *Chlorella* sp. (Franklin 2000; Charles 2002), *C. reinhardtii* (Lavoie 2014), or *Ankistrodesmus* sp. (Cheng 2023). Since U toxicity has been studied in media with different U speciation and bioavailability, a comparison of U tolerance between these microalgae is not relevant. In this study, we compared the tolerance of *Coelastrella* sp. PCV, *C. reinhardtii*, and *C. vulgaris* in similar conditions and could determine that *Coelastrella* sp. PCV is the more U-tolerant species. In the TAP LoP medium, *C. reinhardtii* and *C. vulgaris* cells died at 100 µM uranyl nitrate while *Coelastrella* sp. PCV was still able to grown (although at a reduced rate) and maintain fully efficient photosynthesis up to 200 µM uranyl nitrate (**Figure 3**).

A U-tolerant microalgal species has been formerly isolated from open pits and tailings ponds at Spanish mining sites extensively contaminated with metals, primarily U and zinc. (Baselga-Cervera 2018). This *Chlamydomonas* sp. has the ability to grow rapidly in natural contaminated waters from U mine and in synthetic media containing U. A comparison of U tolerance between this *Chlamydomonas* sp. and *Coelastrella* sp. PCV is not possible for the above mentioned reasons. However, it can be hypothesized that these two U-tolerant algae are derived from different processes. On the one hand, the capacity of *Coelastrella* sp. PCV to tolerate elevated U concentrations is probably due to its genetic background (prior to colonization of U-containing liquid waste) and some physiological and molecular acclimation to the U-contaminated environment from which it was isolated. Indeed, the short duration (a few weeks) and low intensity (1.2 ppm U) of the stress cannot be considered an extreme condition leading to genetic adaptation through the selection of spontaneous mutations (Garcia-Balboa 2013). On the other hand, the U-tolerant *Chlamydomonas* sp. is resulting from an adaptive process that took place over several years to ensure survival under very hostile conditions (U ranging from 25 to 48 ppm) (Baselga-Cervera 2018).

Several mechanisms of tolerance to U have been described in microorganisms (Kolhe 2018). They include metal biosorption onto the cell wall, metal uptake and then sequestration by compartmentalization, metal detoxification through binding to chelators (metabolites or proteins/polypeptides), metal exclusion by efflux transport, metal biomineralization (formation of an insoluble mineral precipitate inside or outside the cell), or metal bioreduction (reduction of soluble U(VI) into insoluble U(IV)). When challenged with U, *Coelastrella* cells very rapidly capture the metal from the medium (adsorption and absorption) and then progressively release U into the environment during prolonged exposure (**Figure 4**). Thus, *Coelastrella* sp. PCV behaves as an excluder to limit the toxic outcomes of U. The process behind U efflux transport by *Coelastrella* is still unknown. However, it is only active when cells are healthy and metabolically active (efflux was not observed in cells exposed to 400 µM U in which photosynthesis is significantly reduced, **Figure 2B-C**) but not when the amount of U accumulated in cells is too low to trigger a defense response (as it is the for natural polluted waters from the Rophin wetland, **Figure 7**). Vogel *et al*. (2010) previously observed desorption of U from *C. vulgaris* living cells during prolonged exposure (2-3 days) to the metal. The authors suggested, but did not demonstrate, that the release of metabolites such as organic acids could explain this behavior. Recently, an unusual U biomineralization process was described in a microalga of the genus *Ankistrodesmus* (Cheng 2023). While the mechanisms of U biomineralization described in some bacteria and fungi involve the formation of U-phosphate complexes (Kolhe 2018), intra- and extracellular crystals of a mineral containing U and potassium (compreignacite) were observed in *Ankistrodesmus* sp., suggesting an important role of potassium in response to U stress. Further analyses will be required to specifically address whether U biomineralization is part of the defense mechanisms evolved by *Coelastrella* sp. PCV to tolerate U.

The rapid, reversible and energy-independent biosorption of U onto various microalgae has been described (Horikoshi 1979; Zhang 1997; Fortin 2004; Vogel 2010; Baselga-Cervera 2018). It is difficult to compare U biosorption capacity in different species because experimental parameters such as algal biomass, medium composition, pH, contact time or U concentration that influence this process are not similar. However, our data showed that the ability of *Coelastrella* sp. PCV to capture U is remarkably high. Indeed, titration of U accumulation indicated that up to 600 mg U per g DW was sorbed to *Coelastrella* cells, of which 240 mg.g^−1^ DW was tightly bound or internalized (i.e. not extractable with carbonate). These values do not reflect the maximum accumulation capacity of *Coelastrella* sp. PCV because titration of this process did not show saturation, even at 400 µM U (**Figure 5**). For comparison, the original and artificially modified U-tolerant *Chlamydomonas* sp. isolated from U mine tailings accumulated 4 to 6 mg U per g DW (Baselga-Cervera 2018) and the highest adsorption capacity reported to date for a green microalga was 75 mg.g^−1^ DW for *Scenedesmus obliquus* (Zhang 1997), a species belonging to the Scenedesmaceae family, as does *Coelastrella* sp. PCV. It is well known that the capacity of algae to bind cationic metals is due to the anionic nature of functional groups of cell wall components (Spain 2021). The involvement of carboxylic and organic/inorganic phosphate groups in the coordination of U by *C. vulgaris* cells was demonstrated (Gunther 2008; Vogel 2010). The very high capacity of *Coelastrella* sp. PCV to accumulate U suggests i/ a unique composition of its cell wall as compared with other microalgae, and/or ii/ a high potential to absorb the radionuclide inside algal cells. Cell wall analysis of *Coelastrella* sp. (3-4) revealed a high quantity of polysaccharides, and thus a highly hydrophilic cell surface, and pointed out important changes in cell wall composition and thickness in relation with growth conditions (Spain 2022). The U uptake mechanisms have been recently characterized in *Saccharomyces cerevisiae* (Revel 2022) but are still unknown in green microalgae. Deciphering the composition and dynamics of the cell wall during U stress, together with U uptake mechanisms, will deserve special attention to understand the unique ability of *Coelastrella* sp. PCV to capture U.

Besides its ability to tolerate a high concentration of U in a synthetic culture medium, *Coelastrella* sp. PCV was able to grow and maintain high photosynthetic activity in natural waters from a wetland near a reclaimed U mine (**Figure 6**). The amount of U found in waters from Rophin is similar to that reported in natural waters from U mines in Germany (Bernhard 1998) but lower than that of a U mine tailings pond where a U-tolerant *Chlamydomonas* species was isolated (Baselga-Cervera 2018). The concentration of essential elements in waters from Rophin is two to three orders of magnitude lower than in artificial algal culture media and the concentration of TOC ranges from 4 to 19 mg.mL^−1^ (**Table 1**). Thus, the speciation, bioavailability and toxicity of U are different in natural or synthetic media used in this study (Markich 2002). In a single one-week growth cycle, *Coelastrella* was able to capture 25 to 55% of U from natural contaminated waters (**Figure 7**). The efficiency of U remediation was not related to the initial U level but was inversely correlated with the concentration of TOC (**Table 1**), as suggested by the decreased bioavailability of U with increasing TOC level (Hogan 2005, Trenfield 2011; Trenfield 2012). The fact that *Coelastrella* sp. PCV was able to remediate waters containing nanomolar amount of U is another illustration of the very high intrinsic capacity of the alga to capture U, even when metal speciation is complex (Markich 2002). In addition to U, *Coelastrella* sp. PCV was able to remediate part of lead contaminating waters from Rophin. This suggests that *Coelastrella* sp. PCV has the potential to successfully address remediation of different contaminating metals, as previously described for the *Coelastrella* sp. (3-4) and *Coelastrella* sp. BGV strains (Plohn 2021; Karcheva 2022).

Due to nutritional deficiency and/or metal toxicity (e.g. U, lead, aluminum), *Coelastrella* sp. PCV accumulated lipid droplets when grown in natural waters from Rophin (**Figure 8**). *Coelastrella* sp. of diverse origins have been shown to accumulate large amounts of neutral lipids (17-22% of total biomass) when cultivated in stress conditions (nutrient deprivation, high salinity, light or temperature) (Nayana 2022). Several *Coelastrella* species have also been reported to produce carotenoids in stress conditions, including *C. astaxantica*, *C. rubescens* or *C. aeroterrestrica* (Nayana 2022). *Coelastrella* sp. PCV has been cultivated in different stress conditions, including nutrient deprivation or metal stress (U and other metals), but none of these treatments resulted in carotenoid accumulation. This suggests that metabolic changes induced by nutrient or metal stress are not similar in all *Coelastrella* species (Karpagam 2018).

## 5. Conclusions

In this study, we characterized a resilient microalgal species demonstrating both tolerance to high U concentration in artificial culture media and the ability to survive in nutrient-deprived natural waters containing substantial amounts of toxic elements (including U, lead or aluminum). As it is much more tolerant to U than *C. reinhardtii* or *C. vulgaris*, *Coelastrella* sp. PCV is a very promising model for identifying mechanisms of response and adaptation to U stress, in particular the efflux process that probably protects algal cells from the toxic effects of U during a prolonged stress period. To support this assumption, the characterization of (hyper)tolerant microbial species has allowed the identification of novel mechanisms of U tolerance, including biomineralization processes or a high-affinity U-binding protein (e.g. Choudhary 2011, Pinel-Cabello 2021, Gallois 2022). *Coelastrella* sp. PCV also demonstrated very interesting properties for treating natural or wastewaters with low to high U contamination. On the one hand, this fast growing and high biomass-producing microalga could be used to reclaim heavily contaminated environments, taking advantage of its remarkable ability to capture U. On the other hand, *Coelastrella* sp. PCV could be grown in natural softwaters contaminated with moderate amounts of toxic metals, allowing both remediation of the habitat and valorization of the algal biomass that accumulates lipids.

## Declaration of Competing Interest

The authors declare that they have no known competing financial interests or personal relationships that could have appeared to influence the work reported in this paper.

## CRediT authorship contribution statement

Camille Beaulier: conceptualization, investigation, writing – original draft, review and editing. Marie Dannay: conceptualization, investigation. Fabienne Devime: investigation. Célia Baggio: investigation. Nabila El Sakkout: investigation. Camille Raillon: investigation. Olivier Courson: investigation. Jacques Bourguignon: funding acquisition, investigation, writing - review and editing. Claude Alban: conceptualization, investigation, writing - review and editing. Stéphane Ravanel: supervision, funding acquisition, conceptualization, investigation, writing - original draft, review and editing.

## Acknowledgments

This work has received funding from the University Grenoble Alpes (PhD fellowship to Camille Beaulier), the Plant Biology and Breeding division from INRAE (DemoniaCo project), the “Nucléaire, énergie, environnement, déchets, société - NEEDS” program from the CNRS (INSPECT project), the “Diversity of biological mechanisms” program from the Institute for Biological Sciences at CNRS (SilverCoela project), and the Agence Nationale de la Recherche (ANR-21-CE34-0004, DemoniaCo project; ANR-17-EURE-0003, CBH-EUR-GS). We gratefully acknowledge Dr. Jonathan Przybyla-Toscano for critical reading of the manuscript and helpful discussions.

## Supplementary material

**Figure S1:**
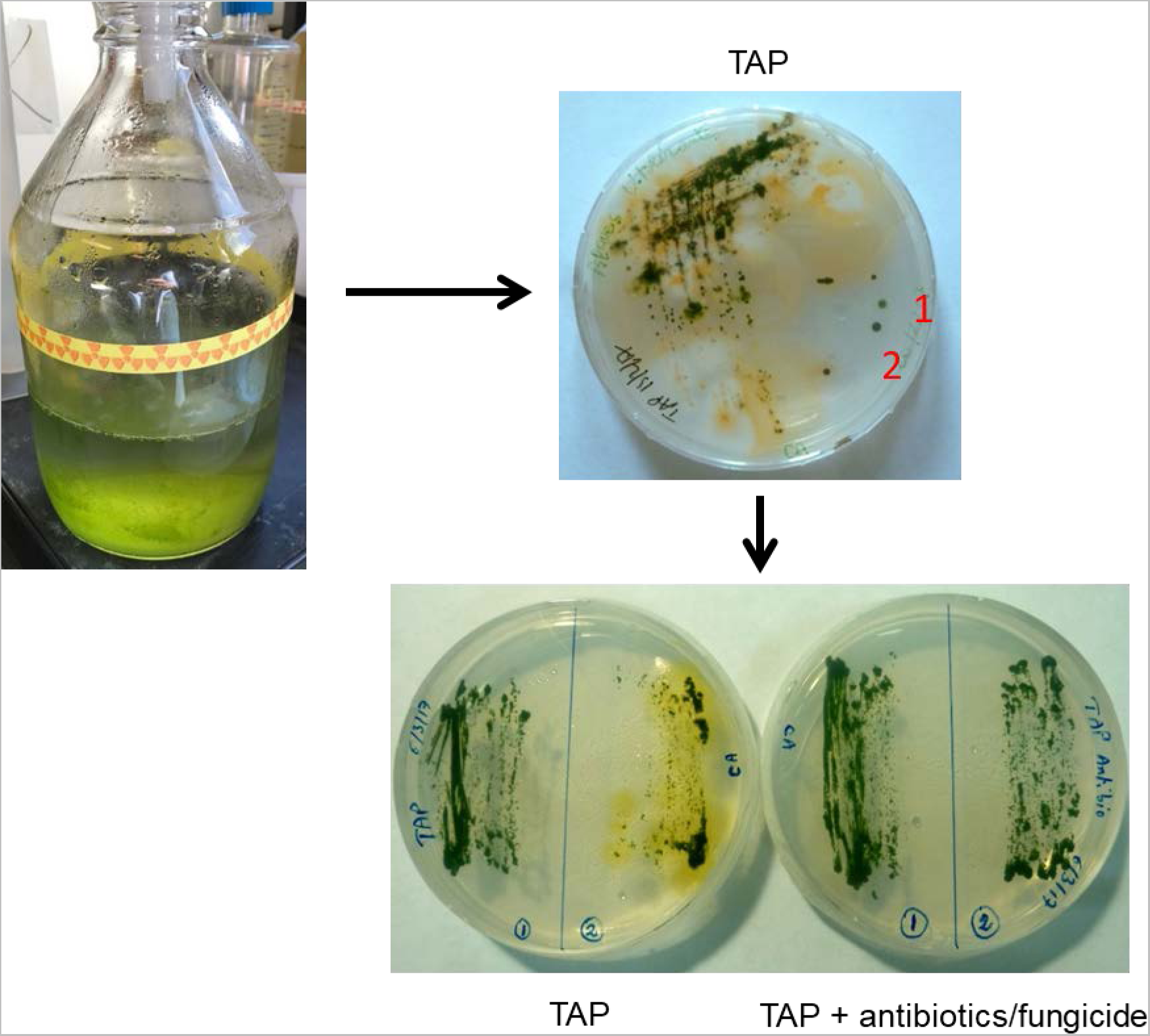
Isolation of *Coelastrella* sp. PCV from liquid wastes contaminated by uranium. The algal bloom developed in laboratory wastes from plant culture medium contaminated with uranyl nitrate (about 5 µM). The original sampling streaked on TAP-agar was contaminated with bacteria and fungi. Single algal colonies (1 and 2) could be cleared of contaminants by two successive rounds of selection on TAP-agar supplemented with antibiotics (ampicillin and cefotaxime) and a broad-spectrum fungicide (carbendazim). Plates were incubated at 21°C in continuous light (20 µE) for 7 days.

**Figure S2:**
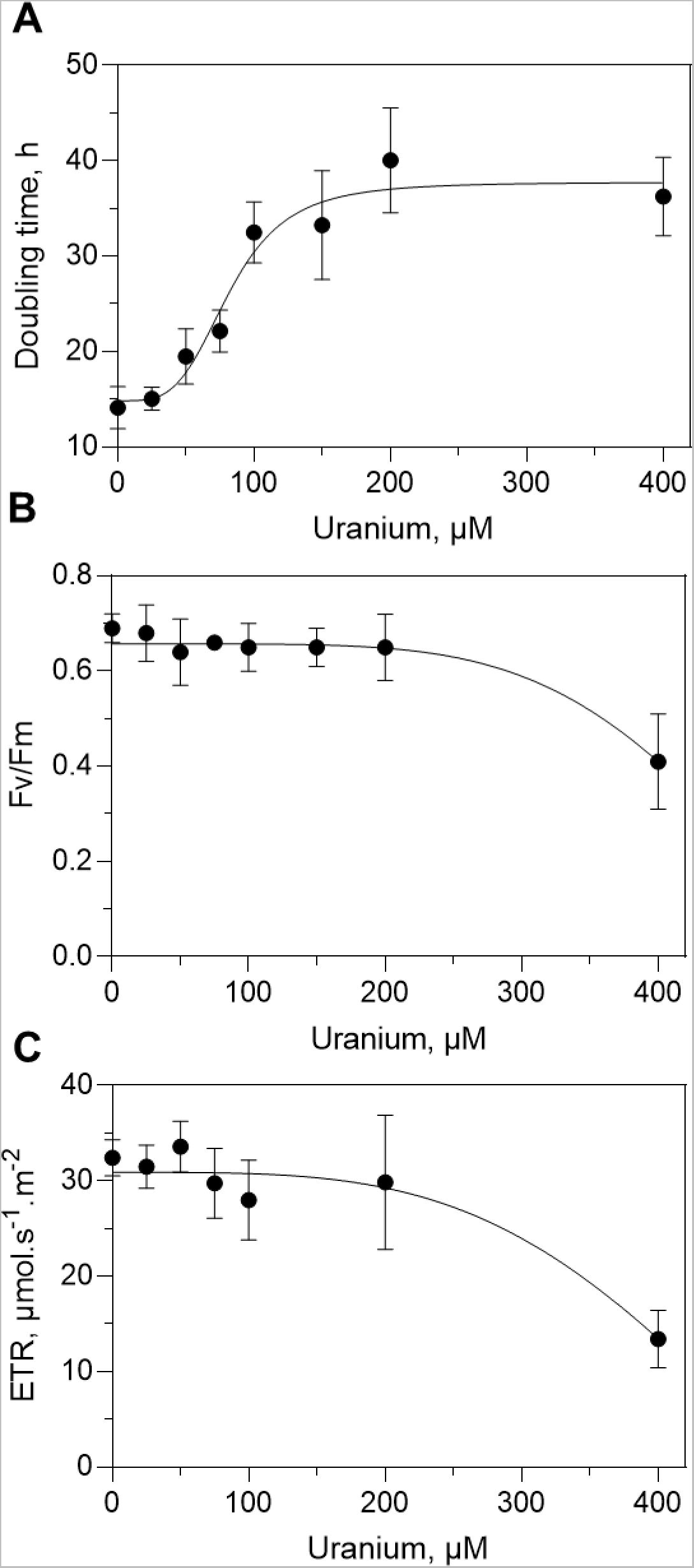
Inhibition of growth and photosynthesis by uranium in *Coelastrella* sp. PCV. A-Effect of U on growth. Doubling times were determined at day 2 from growth curves displayed in Figure 2, plus additional experiments with a similar design (for 25 and 75 µM uranyl nitrate). B,C-Effect of U on photosynthesis. Fv/Fm in dark-adapted cells (B) and electron transfer rates at a light irradiance of 336 µE (C) were measured at day 2 (see Figure 2). Dose-response curves have been drawn from n=5 to 8 independent experiments (mean ± SD) and fitted using sigmoidal equations, indicating IC50 values >80 µM for doubling time, >500 µM for Fv/Fm, and >400 µM for ETR.

## Notes

### Competing Interest Statement

The authors have declared no competing interest.

